# AMPK agonism optimizes the *in vivo* persistence and anti-leukemia efficacy of chimeric antigen receptor T cells

**DOI:** 10.1101/2024.09.26.615290

**Authors:** Erica L Braverman, Mengtao Qin, Herbert Schuler, Harrison Brown, Christopher Wittmann, Archana Ramgopal, Felicia Kemp, Steven J Mullet, Aaron Yang, Amanda C Poholek, Stacy L Gelhaus, Craig A. Byersdorfer

## Abstract

**BACKGROUND:** Chimeric antigen receptor T cell (CART) therapy has seen great clinical success. However, up to 50% of leukemia patients relapse and long-term survivor data indicate that CART cell persistence is key to enforcing relapse-free survival. Unfortunately, ex vivo expansion protocols often drive metabolic and functional exhaustion, reducing in vivo efficacy. Preclinical models have demonstrated that redirecting metabolism ex vivo can improve in vivo T cell function and we hypothesized that exposure to an agonist targeting the metabolic regulator AMP-activated protein kinase (AMPK), would create CARTs capable of both efficient leukemia clearance and increased in vivo persistence.

**METHODS:** CART cells were generated from healthy human via lentiviral transduction. Following activation, cells were exposed to either Compound 991 or DMSO for 96 hours, followed by a 48-hour washout. During and after agonist treatment, T cells were harvested for metabolic and functional assessments. To test in vivo efficacy, immunodeficient mice were injected with luciferase+ NALM6 leukemia cells, followed one week later by either 991- or DMSO-expanded CARTs. Leukemia burden and anti-leukemia efficacy was assessed via radiance imaging and overall survival.

**RESULTS:** Human T cells expanded in Compound 991 activated AMPK without limiting cellular expansion and gained both mitochondrial density and improved handling of reactive oxygen species (ROS). Importantly, receipt of 991-exposed CARTs significantly improved in vivo leukemia clearance, prolonged recipient survival, and increased CD4+ T cell yields at early times post-injection. Ex vivo, 991 agonist treatment mimicked nutrient starvation, increased autophagic flux, and promoted generation of mitochondrially-protective metabolites.

**DISCUSSION:** Ex vivo expansion processes are necessary to generate sufficient cell numbers, but often promote sustained activation and differentiation, negatively impacting in vivo persistence and function. Here, we demonstrate that promoting AMPK activity during CART expansion metabolically reprograms cells without limiting T cell yield, enhances in vivo anti-leukemia efficacy, and improves CD4+ in vivo persistence. Importantly, AMPK agonism achieves these results without further modifying the expansion media, changing the CART construct, or genetically altering the cells. Altogether, these data highlight AMPK agonism as a potent and readily translatable approach to improve the metabolic profile and overall efficacy of cancer-targeting T cells.

## INTRODUCTION

Chimeric Antigen Receptor T cell (CART) therapy has had a significant impact on the treatment of relapsed/refractory acute B-cell lymphoblastic leukemia, with more than 90% of treated pediatric patients initially achieving remission [1]. However, despite the success of this adoptive cellular therapy, up to 50% of patients relapse after CART treatment, limiting its utility as a long-term cure [2, 3]. Further, CARTs have seen limited success in other cancers, particularly solid tumors. While the reasons for this limited efficacy are many, one of the most prominent concerns relates to the functional status of the injected CART cells. The ex vivo expansion process drives significant activation and differentiation of CARTs, limiting their ability to form memory populations and negatively impacts their in vivo persistence [4]. This combination results in leukemia relapse and restricted tumor clearance in other cancers [5]. As such, identifying methods to augment the in vivo function and persistence of CARTs has become critical to improving their therapeutic efficacy.

Many interventions have shown promise towards improving CART persistence. On the one hand, generating less differentiated, more memory-like CARTs has seen great effect, achieved by driving expression of memory transcription factors, blocking differentiation pathways, and changing the cytokine milieu of the growth media [6–11]. It has also become clear that targeting CART cell metabolism, for example by specifically augmenting their respiratory capacity, is another method to improve their long-term survival. This reprogramming has been achieved by restricting access to certain nutrients during expansion (essentially enforcing a “starvation” program), blocking specific metabolic pathways, or driving expression of mitochondrially-focused genes to augment mitochondrial health and capacity [4, 12–19]. However, many of these approaches, while necessary to increase functionality, have limited translatability to the clinic. For example, nutrient restriction (i.e., by blocking glycolysis), slows CART proliferation and results in fewer CART cells for in vivo transfer. Further, the need to re-engineer expansion media, by removing or adding specific nutrients, poses its own cost and logistical barriers. Finally, while over- or under-expressing certain metabolic genes in CARTs has been effective in mouse models, this added genetic manipulation raises concerns about the oncologic potential of modified CARTs, slowing their path to translation. Even without these barriers, the existence of such a wide array of modifiable pathways raises the question as to which options will create the optimal CART cell product – and how best to achieve many of those goals simultaneously. Put simply, identifying an optimized strategy that does not require extensive manufacturing changes while simultaneously promoting multiple advantageous pathways, will be paramount to achieving more effective CART therapies.

AMP-activated protein kinase (AMPK) is a heterotrimeric cellular energy sensor upstream of a web of metabolic outputs [20] and is best known for recognizing nutrient restriction and reprogramming cellular metabolism towards catabolic energy generation while reducing anabolic growth. We have previously demonstrated that driving AMPK activity in human T cells augments mitochondrial capacity, memory formation, and inflammatory function [21]. In addition, many targets of AMPK’s metabolic reprogramming have been highlighted as being advantageous during ex vivo CART cell expansion, including nutrient restriction, blockade of the mammalian target of rapamycin (mTOR), counteracting reactive oxygen species (ROS), promotion of mitochondrial biogenesis, and enhancement of autophagy [6, 8, 9, 11, 12, 15–19, 22]. With AMPK upstream of so many beneficial metabolic programs, we hypothesized that facilitating AMPK signaling during ex vivo expansion would create a metabolically optimal CART product.

Detailed below, we highlight the novel use of a direct AMPK agonist, Compound 991, to metabolically re-program human T cells. Exposing T cells to an AMPK agonist which binds directly to the AMPK heterotrimer [23] created metabolically augmented cells with significantly improved in vivo anti-leukemia activity. Interestingly, 991 exposure did not drive memory reprogramming but instead orchestrated a network of metabolic changes including increased autophagic flux, enhanced fatty acid oxidation, and generation of mitochondrially-protective metabolites. Together, these changes created CARTs with improved in vivo persistence, particularly within the CD4+ compartment. In total, these studies highlight the potential for short-term, direct AMPK agonist treatment to rewire the metabolic capacity of CARTs, providing an easily translatable method that simultaneously modifies multiple beneficial pathways to improve CART therapy.

## RESULTS

### 991 treatment facilitates AMPK activity without restricting expansion

We hypothesized that expanding CART cells in the presence of a direct AMPK agonist would metabolically optimize them for in vivo function. To test this hypothesis, we first interrogated whether Compound 991 treatment activated AMPK without restricting growth or viability. Human T cells were isolated and stimulated with anti-CD3/CD8 Dynabeads for 5 days, removed from the beads, and split into control (DMSO) and 991 treated groups. The AMPK heterotrimer is active when the alpha subunit, containing the kinase domain, is phosphorylated on Thr172 [24]. Dosing experiments indicated stable phosphorylation of AMPKα Thr172 for 48 hours following 991 exposure, leading to a final treatment schedule where 991 was added to T cell cultures for two 48 hours cycles (96 hours of total exposure), followed by a 48-hour washout period (Fig1A). Measurement of AMPKα phosphorylation confirmed increased activation of AMPK following 991 treatment (Fig1B), with no significant impact on cell growth or expansion through day 11 (Fig1C). To assess whether equivalent cell numbers indicated similar proliferation or a combination of proliferative differences and a change in cell survival, we assessed T cell proliferation by measuring incorporation of the thymidine analogue Bromodeoxyuridine (BrdU). In line with our expansion data, there was no difference in BrdU uptake on day 9, following 96 hours of agonist treatment on Day 9 (Fig1D). Interestingly, when BrdU incorporation was measured at the end of the culture period on Day 11, there was now a significant increase in the proliferation of 991-treated cells (Fig1E). Given AMPK’s well-documented roles in optimizing metabolic fitness, we hypothesized this ongoing cell turnover was due to enhanced metabolic capacity, which then allowed for a sustained proliferative effort despite the increasing distance from their original stimulation. We therefore sought to measure the impact of 991 exposure on subsequent metabolic reprogramming.

**Figure 1.**
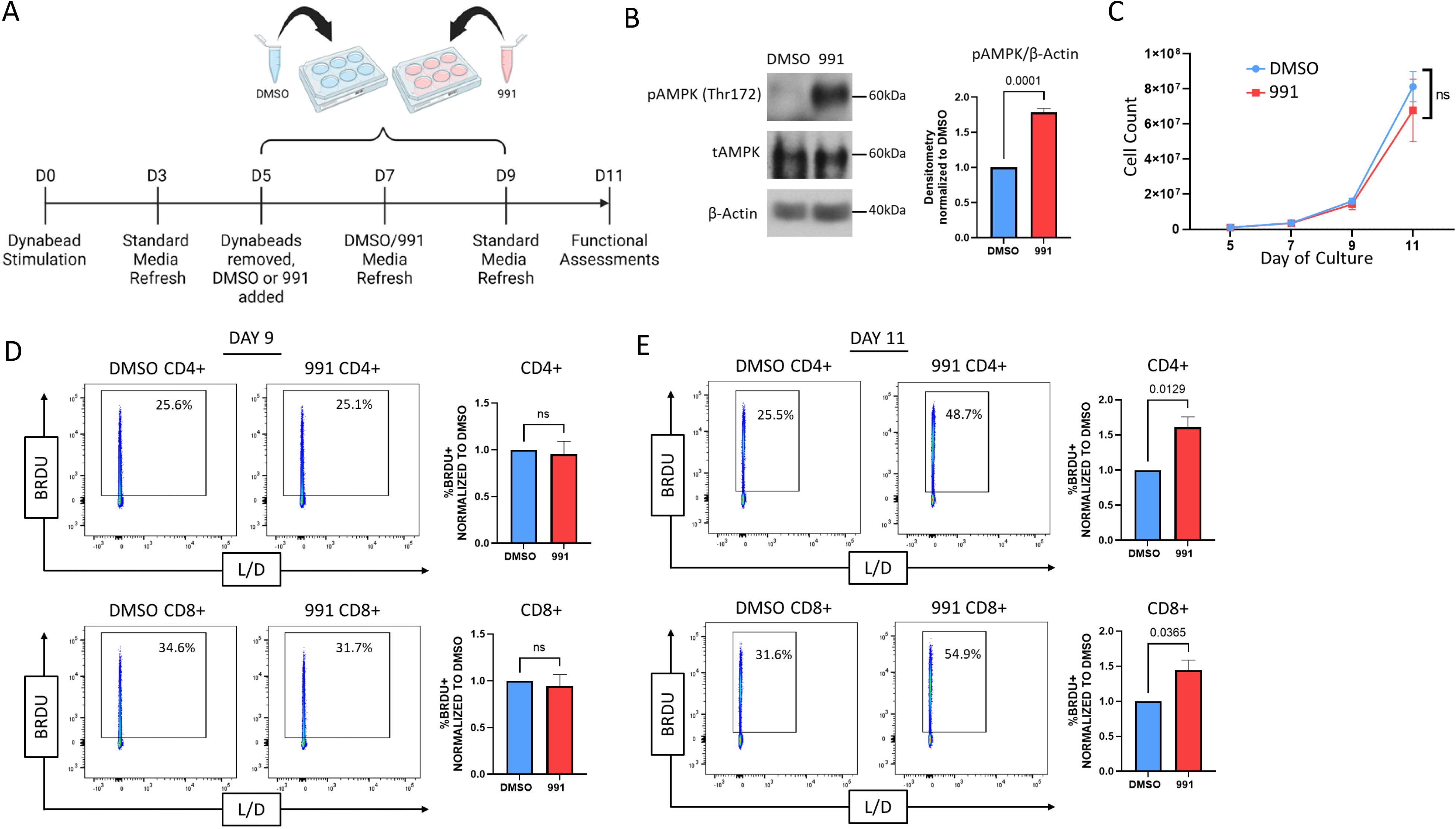
991 treatment drives AMPK activity in Human T cells without restricting expansion. *A,* Schematic of Compound 991 treatment protocol. *B,* Proteins from human T cells treated with 991 or DMSO control were precipitated on Days 7-9 and phosphorylation of AMPKα on Thr172 (to detect AMPK activation) was measured by immunoblot. Accompanying densitometry was quantitated on cells obtained from multiple donors using ImageJ software, followed by normalization of 991-treated levels within each sample to DMSO controls. *C*, Human T cells were manually counted on Days 5, 7, 9, and 11 and counts plotted to demonstrate expansion over time. *D-E*, DMSO and 991-treated cells were incubated with BrdU for 2 hours on Day 9 **(D)** or 11 **(E)** of culture, followed by staining for BrdU incorporation. All data were obtained on 3 or more independent human donor samples. Numbers above the graphs represent statistical significance as determined by paired Student T test.

### 991-treated human T cells gain mitochondrial capacity and efficiency

To gauge the impact of 991 treatment on T cell metabolism, we utilized the Seahorse Metabolic Analyzer Mitostress test to measure mitochondrial capacity. On Day 11 (48 hours post 991 removal), 991-treated cells increased their oxygen consumption rates (OCR) and spare respiratory capacity (SRC) (Fig2A). We hypothesized these increases might be secondary to an increase in total mitochondria, particularly as AMPK is known to activate (PGC1α), a transcription factor responsible for promoting mitochondrial biogenesis. Staining with MitoTracker revealed increased mitochondrial density in 991-treated cells (Fig2B), which correlated with elevated PGC1α expression during 991-treatment (Fig2C). To better understand if these metabolic changes would persist following subsequent stimulation, we restimulated cells on Day 11 and repeated our metabolic assessments (Fig2D). As shown in Fig2E, augmented mitochondrial activity continued following activation, with increases in both OCR and SRC. Of note, driving mitochondrial metabolism can also generate increased levels of reactive oxygen species (ROS), which can be damaging to cells at high levels. Reassuringly, enhanced AMPK signaling improved ROS handling, which we hypothesized was likely contributing to the ability of 991-treated cells to tolerate increased mitochondrial respiration. Consistent with this interpretation, 991-pretreated cells had lower ROS burden following 24 hours of stimulation (Fig2F). We also regularly recovered greater numbers of 991-treated cells following 72 hours of restimulation (Fig2G), consistent with improved ROS handling supporting an increase in cellular proliferation. Altogether, these data demonstrate that 991-treatment facilitates mitochondrial biogenesis and enhances mitochondrial function, allowing for increased metabolic capacity and improved cellular expansion upon *in vitro* re-stimulation.

**Figure 2.**
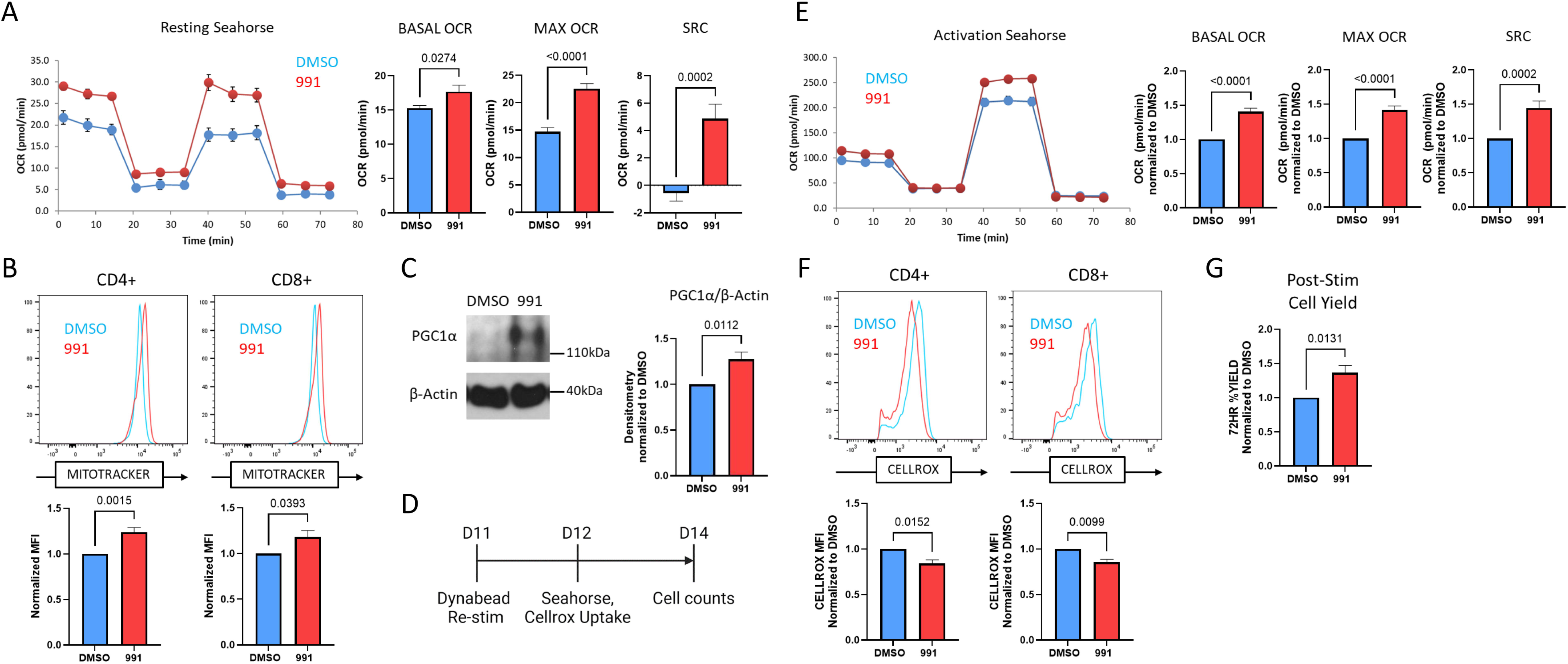
991-treated human T cells gain mitochondrial capacity. *A*, Resting Day 11 DMSO- and 991-treated T cells were assessed for oxidative capacity utilizing the Seahorse Metabolic Analyzer. Bar graphs represent data from 2 individual human donors. *B*, DMSO- and 991-treated T cells were incubated with MitoTracker Green to measure mitochondrial density. Bar graphs represent median fluorescence intensity (MFI) data for 4 human donors. *C*, DMSO- and 991-treated cells were lysed as in Figure 1B and total PGC1a protein measured via immunoblot. Densitometry is shown for 4 human donors. *D*, Re-stimulation schematic. *E*, Human T cells were assessed for oxidative capacity following re-activation with Dynabeads for 24 hours utilizing the Seahorse Metabolic Analyzer. *F*, Re-activated human T cells were incubated with CellROX dye to measure reactive oxygen species. G, Cells were counted at the time of re-stimulation, and again 72 hours later, to determine percentage cell yield. Unless otherwise stated, bar graphs represent composite data from three or more independent human donors. In panels B-G, bar graph data from 991-treated cells was normalized back to DMSO-treated controls.

### AMPK agonist treatment improves CART anti-leukemia activity and prolongs survival in a xenograft model

With data supporting improved metabolic fitness in agonist-treated T cells, we next tested whether 991 pre-treatment improved the function of CART cells targeting leukemia. Human CART cells were generated via lentiviral transduction utilizing a CD19-targeting CAR (Fig3A) and expanded in the presence of the 991 agonist on the same schedule as the polyclonal human T cells in Figures 1 and 2 (Fig3B). We first confirmed that 991-treatment similarly enhanced the metabolic capacity of CART using the Seahorse metabolic analyzer. 991-treated CARTs at rest (Fig3C), as well as those following overnight activation with CD19+ NALM6 leukemia cells (Fig3D), enhanced their mitochondrial capacity. To measure in vivo CAR T cell efficacy, we transferred luciferase expressing NALM6 cells into immunodeficient NSG mice followed one week later by 3e6 CART cells (Fig3E). Standard CART cells transferred into NALM6-bearing NSG mice delayed leukemia growth compared to the leukemia-only control. However, all DMSO-treated CART cell recipients eventually succumbed to lethal leukemia. In sharp contrast, 991-treated CARTs dramatically improved leukemia control, with 54% of 991-treated CART recipients (6/11) remaining leukemia-free through the end of the experiment (Fig3F). This improved leukemia clearance led to a significant and reproducible improvement in recipient survival, with 73% of mice receiving 991-treated CART cells surviving until day 70 (Fig3G). Together, these data highlight that expanding human CARTs in the presence of the AMPK agonist, Compound 991, creates a superior CART cell product, with a striking improvement in overall leukemia clearance and subsequent recipient survival in our preclinical model.

**Figure 3.**
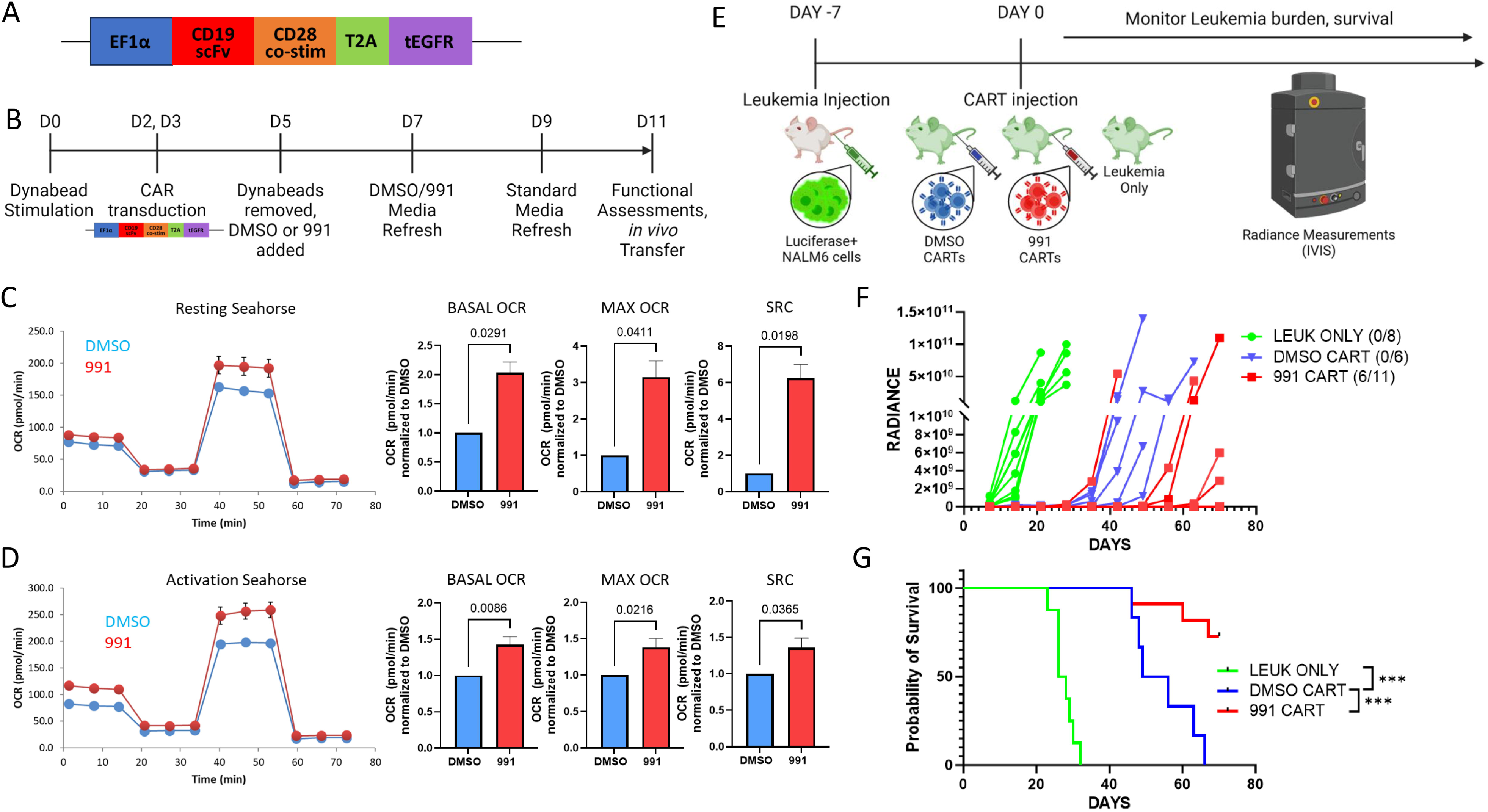
AMPK agonist pre-treatment improves CART anti-leukemic activity and recipient survival. *A-B*, Schematic of CAR plasmid (**A**) and CART transduction and agonist treatment protocol (**B**). *C*, Resting Day 11 DMSO- and 991-treated CART cells were assessed for oxidative capacity utilizing the Seahorse Metabolic Analyzer. Bar graphs represent data from 3 individual human donors. *D*, Human CART cells were re-activated with NALM6 leukemia targets for 24 hours, followed by further assessment of oxidative capacity. Bar graphs represent data from 4 individual human donors and in (**C-D**) are normalized back to DMSO-treated controls. *E*, Schematic of Nalm6 xenograft leukemia model. *F-G*, Radiance measurements (**F**) and survival (**G**) in recipient of Leukemia only (green), DMSO-CART cells (blue), and 991-CART cells (red). n = 8 Leukemia only recipients, 6 DMSO CART recipients, and 11 991-treated CART recipients. Numbers in parentheses (6/11) denote the number of mice remaining leukemia-free over total mice injected. Data are representative of two separate experiments. ***p<0.001, with survival differences between recipients of Leukemia only and 991-treated CART cells being p<0.0001 (not shown).

### 991 treatment upregulates cell cycle and metabolic gene sets without inducing changes in memory or activation markers

Multiple groups have demonstrated that CARTs with memory-like phenotypes demonstrate improved anti-leukemia activity in vivo [11, 15]. However, we found no differences in memory phenotype or activation status in our 991-treated cells (SuppFig1A-D). Since AMPK signaling can also impact cellular transcriptomics, including through direct activation of transcription factors as well as downstream influence on histone deacetylases, we pursued bulk RNA sequencing of Day 11 DMSO- versus 991-treated human T cells. Only a handful of transcripts were significantly altered in either CD4+ and CD8+ T cells, using a p value of <0.05 and log2-fold change of 0.6 (Fig4A-B). However, gene set enrichment analysis (GSEA) uncovered multiple upregulated pathways, with the highest enrichment scores in both CD4 and CD8 T cells clustering within cell proliferation and cell cycle pathways (Fig4C-D), consistent with the higher proliferative rates observed at the end of in vitro culture (Fig1E). The second most enriched gene sets highlighted metabolic pathways (Fig4E-F), with pathways directly related to supporting increased proliferation, including pyrimidine and folate metabolism, as well as a notable enrichment of oxidative phosphorylation and fatty acid oxidation. These latter data are again consistent with the increased oxidative capacity of 991-treated cells (Fig2A-C) and suggest as we hypothesized that increases in cell cycle may result from the enhanced metabolic capacity of 991-treated cells. We also hypothesized that metabolic rewiring downstream of AMPK was likely responsible for the functional advantage of 991-treated CARTs in vivo and therefore sought to understand mechanistically how AMPK was directing metabolism to achieve such impressive results.

**Figure 4.**
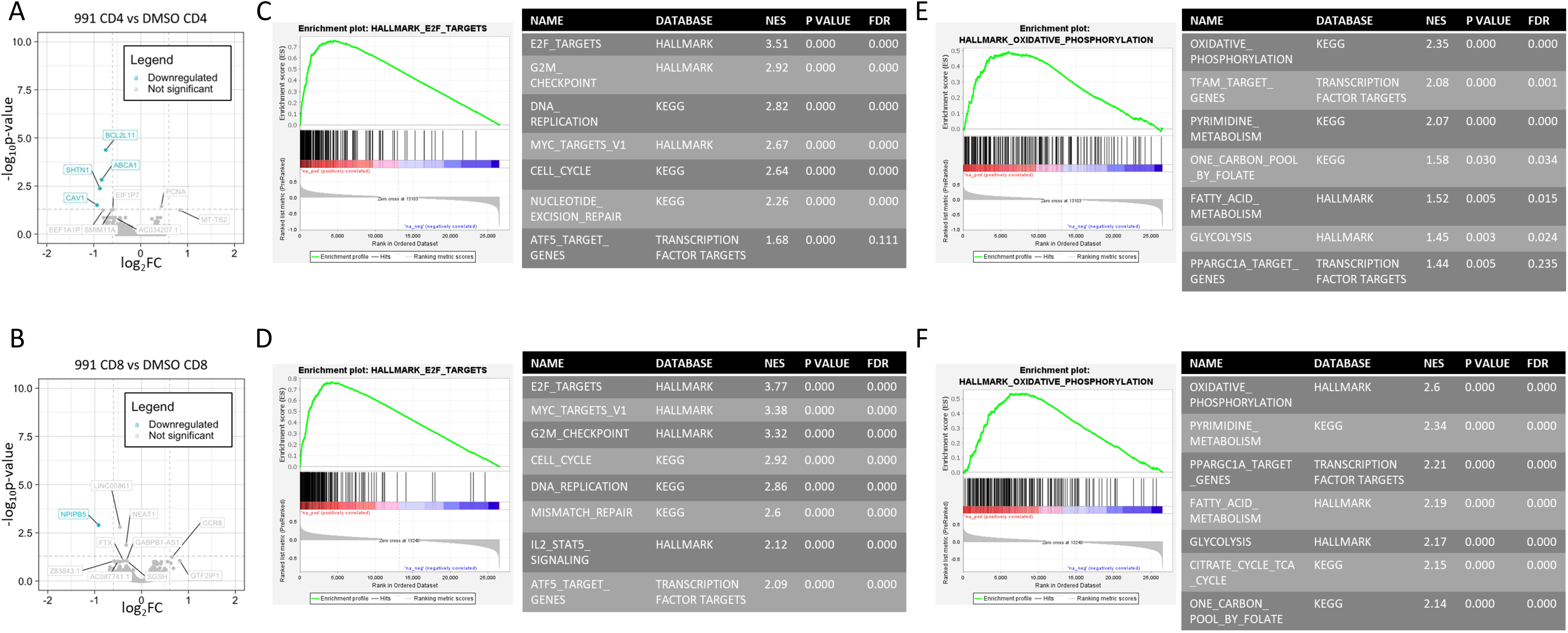
991-treated T cell transcripts are enriched for cell cycle and metabolic gene sets. RNA was harvested from Day 11 T cells from three individual human donors and analyzed for gene expression differences by RNA sequencing. *A-B*, Transcript differences were plotted via log fold-change versus negative log P value, with data points meeting statistical significance highlighted in blue for CD4s (**A**) and CD8s (**B**). *C-F*, Gene sets were then ranked and GSEA performed using comparison to Hallmark, KEGG, and transcription factor databases through the GSEA software (see methods for further details). The highest ranked gene sets were in cell cycling (**C, D**) and metabolism (**E, F**), shown for CD4s and CD8s, respectively. Accompanying tables list additionally enriched cell cycle and metabolic gene sets.

### AMPK agonism simultaneously drives fatty acid oxidation while promoting generation of mitochondrially-protective metabolites

AMPK is well-known for its role in supporting fatty acid oxidation (FAO) and long-chain fatty acids (LC-FAs) can bind directly to AMPK to facilitate its activity. Notably, these LC-FAs use the same binding site as Compound 991 [25]. We therefore hypothesized that T cells treated with 991 would increase their utilization of FAO. Using the oxidation-sensitive dye FAO-blue, we observed increased FAO activity in agonist-treated T cells (Fig5A). This upregulation correlated with a higher sensitivity to etomoxir, the carnitine palmitoyltransferase 1A (CPT1A) and FAO inhibitor, which was read out by a greater reduction in protein translation following etomoxir treatment of agonist-treated cells (Fig5B). We next stained cells for lipid droplets, which serve as storage depots for FA intermediates like triacyl glycerides, before being broken down into single-chain FAs for FAO. In line with an increase in FAO, agonist-treated cells also demonstrated reduced staining with the lipid sensitive dye Nile Red, indicating decreased lipid reserves in agonist-treated cells (Fig5C). Agonist-treated cells also increased expression of CPT1A (Fig5D), the enzyme which facilitates transport of LC-FAs into the mitochondria for subsequent beta oxidation. And despite such significantly upregulated FAO, there was no difference in ROS generation during agonist treatment as measured by CellROX staining (SuppFig2A).

**Figure 5.**
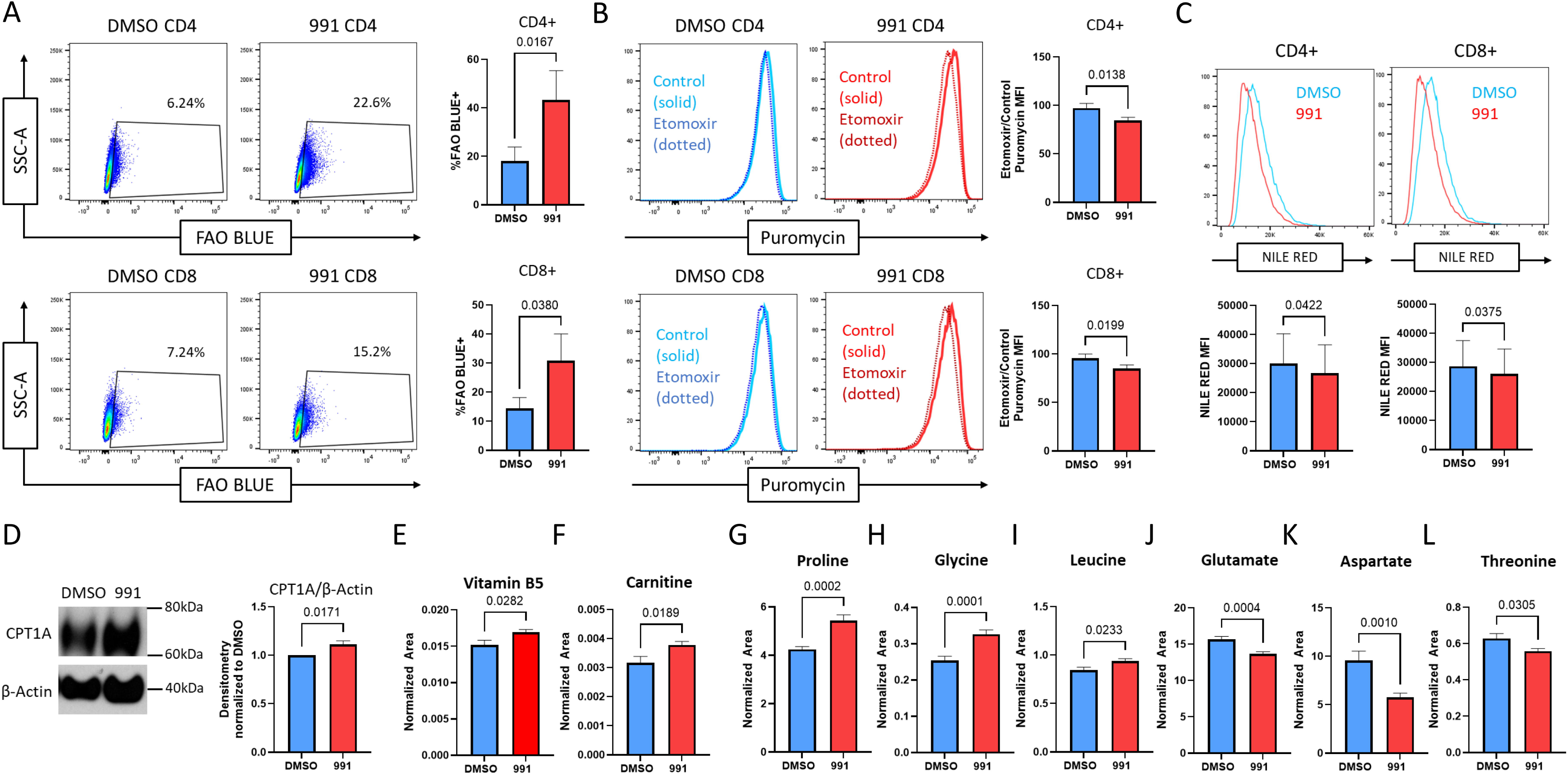
AMPK agonism drives fatty acid oxidation and promotes generation of mitochondrially-protective metabolites. *A*, Cells were incubated with FAOBlue dye for 2 hours, followed by flow cytometry analysis. *B*, Cells were pre-incubated +/− etomoxir, then incubated with puromycin for 30 minutes, followed by staining for puromycin incorporation. Bar graphs represent the MFI of etomoxir-treated group divided by the MFI of the control group for both DMSO and 991-treated cultures. *C*, Cells were incubated with Nile Red dye for 10 minutes followed by flow cytometry analysis. *D*, Total CPT1A protein was measured by immunoblot and densitometry normalized in each sample to DMSO controls. *E-F*, Vitamin B5 (**E**) and free carnitine (**F**) levels were measured by mass spectrometry. *G-L*, Mass spectrometry measured levels of intracellular proline (**G**), glycine (**H**), leucine (**I**), glutamate (**J**), aspartate (**K**), and threonine (**L**). All bar graphs represent data from 3 or more human donors.

Mass spectrometry analysis of intracellular metabolites in 991-treated day 9 cells further identified increased abundance of Vitamin B5 (Fig5E) and carnitine (Fig5F), two additional intermediates necessary for generating fatty-acyl-coA moieties and transporting them across the mitochondrial membrane, respectively. Further inspection of the metabolite data also noted upregulation of multiple amino acids (AAs) known to play a role in mitochondrial health and fitness, including proline, glycine, and leucine (Fig5G-I) [26–30]. Precursors of these AAs, such as glutamate, aspartate, and threonine, were conversely decreased (Fig5J-L), suggesting that AMPK specifically directs production of mitochondrially protective AAs. Altogether, these metabolic data highlight AMPK’s roles, not only in promoting FAO, but also in augmenting production of metabolites with roles in maintaining mitochondrial health and function to support the desired metabolic programming.

### AMPK agonism mimics cellular starvation and upregulates autophagy to enhance metabolic fitness

Some of the earliest literature aimed at improving T cell fitness highlighted the utility of blocking glycolysis during cellular expansion [4]. With GSEA also highlighting enriched glycolytic datasets in 991-treated cells (Fig4E-F), we next sought to understand the role of glycolysis downstream of AMPK agonism. Since AMPK is known to promote glucose uptake, we first quantified the amount of glucose remaining in the media after 48 hours of culture in the presence of 991. In contrast to an expected AMPK-mediated increase in glucose uptake, we regularly found more glucose remaining in the media of 991-treated cultures than in DMSO-treated controls (Fig6A). Supporting this lack of glycolytic activity, media from 991-treated cultures also exhibited reduced lactate content (Fig6B), with a reduction in intracellular hexoses in 991-treated cells (Fig6C). However, the full intracellular metabolite analysis painted a different picture, with marked elevated levels of intracellular lactate (Fig6D). Combined with reduced lactate in the media, these data suggest 991-treated cells are continuing to undergo glycolysis, but are retaining the generated lactate intracellularly instead of secreting it. Interestingly, lactate build-up itself has been demonstrated to reduce cellular glucose uptake [31], which may explain the lack of an expected increase in glucose uptake following agonist treatment. With glycolysis ongoing, despite reduced glucose uptake, we sought to understand where cells were sourcing their sugar carbons. To do this, we performed pathway analysis on untargeted metabolite data to delineate pathway changes in our cells. Interestingly, the top four most significantly upregulated metabolic pathways following 991 agonist treatment concerned the breakdown of alternative sugar sources, including glycogen (Fig 6E-F). Pathways involving nucleotide metabolism and mitochondrial performance were also highlighted, again supporting our GSEA results and the observed metabolic activity of agonist-treated cells. Further, an increased reliance on intracellular sugar breakdown, alongside lactate retention, are both in line with cells exhibiting a nutrient starvation response. If enhanced AMPK signaling were indeed promoting a starvation response, we would expect to find an increase in autophagic flux, as cells looked for a way to break down additional energy sources. Such a finding would be of particular interest since increased autophagy, in the setting of nutrient restriction, enforces metabolic efficiency in T cells during in vitro expansion [32].

**Figure 6.**
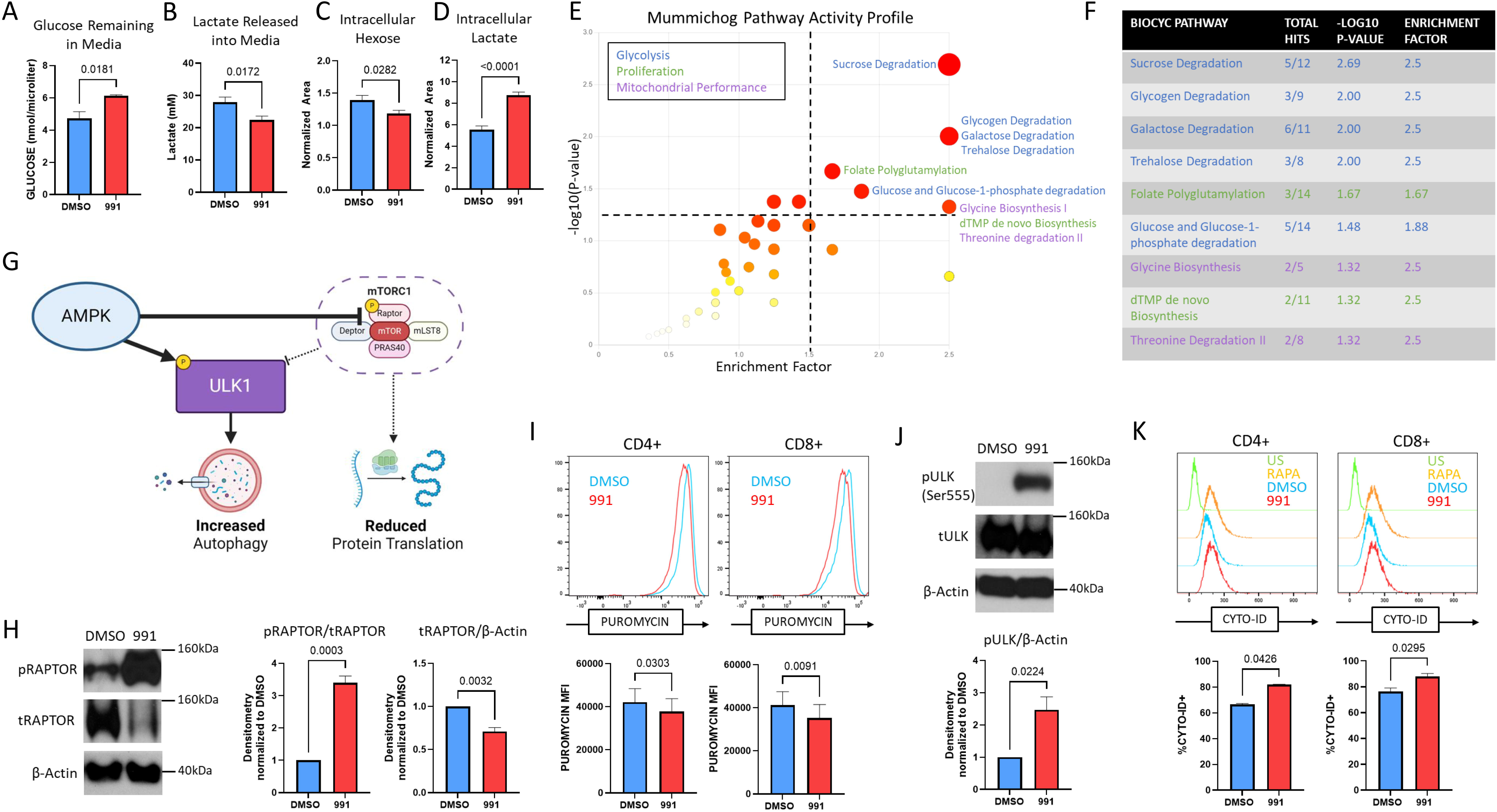
AMPK agonism mimics cellular starvation. *A-B*, Media recovered from 48-hour cultures (+/− 991) was assessed for total glucose (**A**) and lactate (**B**) levels. *C-D*, Intracellular hexose (**C**) and lactate (**D**) content was measured by mass spectrometry in T cells on day 9 of culture. *E*-*F*, Untargeted metabolite data analyzed using Metaboanalyst software. Pathways with an enrichment factor >1.5 are highlighted. *G*, Proposed interactions between AMPK, mTOR, and ULK1. *H*, Immunoblot for phosphorylated and total Raptor levels on Days 7-9 of treatment. Bar graphs represent data from multiple donors, with 991-treated results normalized to DMSO controls. *I*, Cells were incubated with puromycin for 2 hours, followed by intracellular staining for puromycin incorporation. *J*, Immunoblot for phosphorylated ULK1 protein in day 9 cells, with values from multiple donors normalized to DMSO controls. *K*, Day 7 cells +/− 991 were incubated with CYTO-ID dye for 30 minutes and incorporation assessed by flow cytometry. Incubation with rapamycin served as a positive control. Bar graphs represent values from three human donors, except for the CYTO-ID data shown in K, which was two donors.

Although retention of lactate itself promotes cellular autophagy [33], AMPK is also a well-known driver of autophagy, both by activating Unc-51 like kinase 1 (ULK1) and by restricting mTOR-mediated ULK inhibition through phosphorylation of the mTORc1 complex protein, Raptor (Fig6G). We first tested for mTOR inhibition by measuring total and phosphorylated Raptor levels, noting that a large role for AMPK is to target Raptor for phosphorylation-dependent degradation [34]. In 991-treated cells, phosphorylation of Raptor was significantly increased while total Raptor protein levels were significantly decreased (Fig6H). Without a fully functional mTORc1 complex to signal the availability of amino acids, 991-treated cells regularly reduced their translational activity in line with their sense of lower amino acid levels (Fig6I) [35]. Such a reduction in protein translation has independently been highlighted as a further mechanism to improve in vivo T cell function [36]. Meanwhile, 991-treatment increased phosphorylation of ULK1 (Fig6J), concomitant with an increase in cellular autophagy (Fig6K). Together, these data suggest that AMPK agonist treatment drives cellular programming reminiscent of the response to nutrient starvation, increasing availability of intracellular energy sources through autophagy while reducing high energy expenditure by decreasing protein translation.

### Improved leukemia control correlates with increased survival of 991-treated CD4+ CART cells

Our data suggest that AMPK agonism metabolically reprograms cells towards pathways which facilitate cellular fitness. We therefore hypothesized that the mechanism of improved leukemia clearance and subsequent improved survival in our pre-clinical model could be either enhanced initial CART expansion and/or prolonged in vivo persistence of the CARTs over time. Importantly, recent CART clinical data has highlighted the importance of CART cell persistence, particularly within the CD4+ compartment, to mediate effective long-term, leukemia-free survival [3]. To investigate the etiology of improved leukemia clearance, we repeated our leukemia dosing but sacrificed a cohort of mice at either Day 3 or Day 5-7 (one week) post-CART injection (Fig7A). Recipients were injected with BrdU just prior to harvest to measure the active proliferation of previously transferred CART cells. We also enumerated T cells from the bone marrow, where NALM6 leukemia cells first expand, as well as the spleen. There were no differences in proliferation of 991-treated CART cells from the bone marrow at either timepoint (Fig7B), but we did note a transient proliferative increase in DMSO CART cells in recipient spleens on Day 3 that was gone by one week (Fig7C). Importantly, recipients of either DMSO- or 991-treated CART cells demonstrated no evidence of active leukemia in the bone marrow at either day 3 or one week (SuppFig3A). Despite increased BrdU positivity in splenic DMSO CARTs on Day 3, there was no difference in total DMSO-treated T cell numbers in the spleen or bone marrow at this time or at one week (Fig7D). In contrast, there was a significant increase in CD4+ 991-treated CART cells in both the bone marrow and spleen by one week (Fig7E) and this CD4+ T cell advantage drove a notable elevation in the CD4/CD8 ratio, which increased further at the two-week time point (Fig7F). Importantly, there were no pre-injection differences in the CD4/CD8 ratios of DMSO- versus 991-treated CART products (SuppFig4A). Together, these data suggest that the increased leukemia control and subsequent effective survival in mice receiving 991 CART cells correlates with improved persistence of CART cells within the CD4+ compartment.

**Figure 7.**
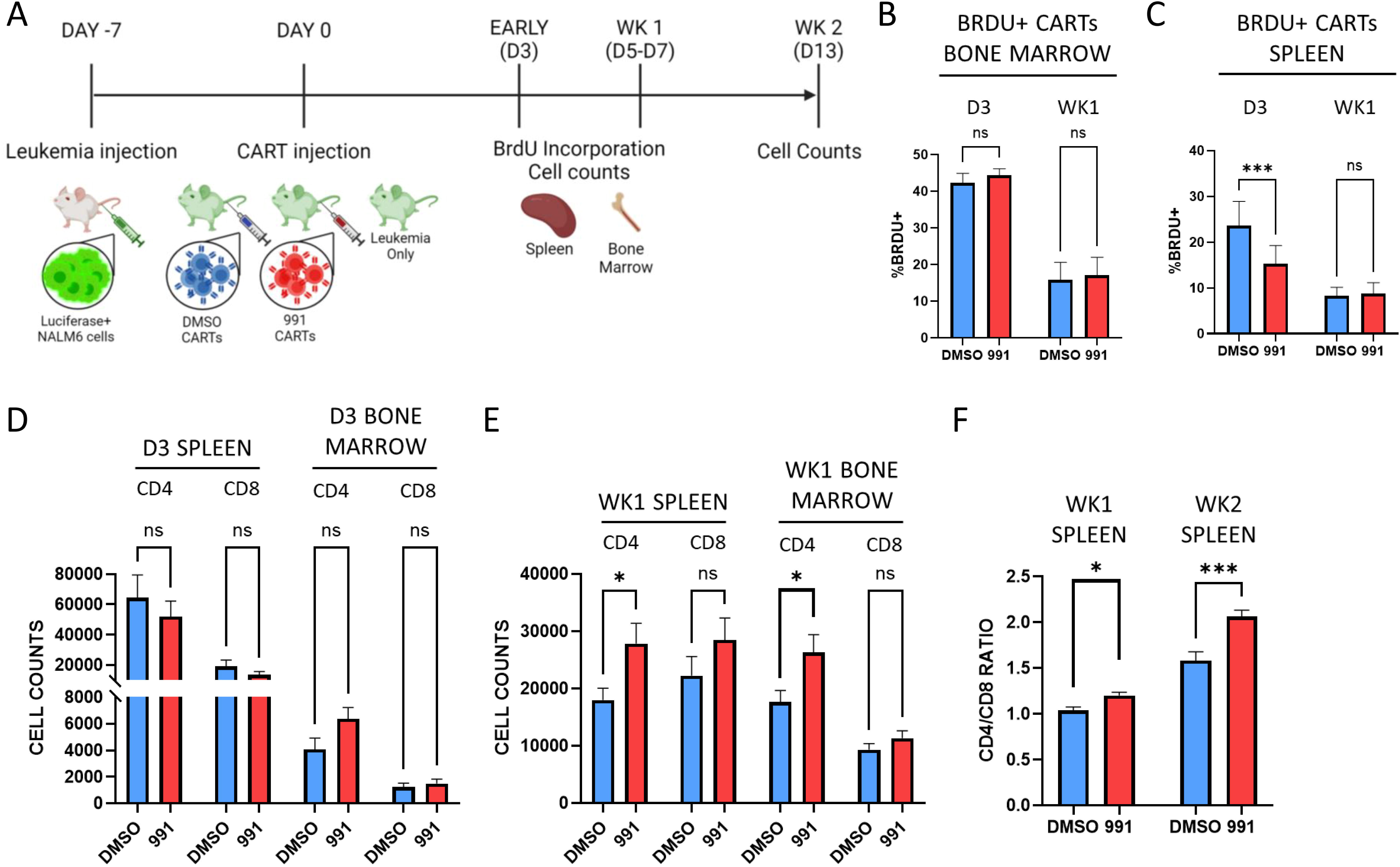
Improved leukemia control correlates with increased numbers of 991-treated CD4+ CART cells. *A*, Timeline of evaluations in our xenograft leukemia model. *B-C*, Mice were injected intraperitoneally with BrdU, followed by spleen and bone marrow harvest 30-60 minutes later. BrdU incorporation was compared between DMSO- and 991-treated CAR T cells in the bone marrow (**B**) and the spleen (**C**), both on Day 3 and up to one-week post-transfer. *D-E*, Total human CD4+ and CD8+ T cell counts were obtained from the spleen and bone marrow on Day 3 (**D**) and after one week (**E**). *F*, CD4/CD8 ratios were calculated in the spleen at one- and two-weeks post-injection. n=7 for both groups in the day 3 bone marrow samples (panels B and D), and n=8 for both day 3 spleen samples (C and D). n=10 mice in both groups at one-week post-injection (C, E, and F) and n=6 and 11 for the week 2 DMSO and 991-treated samples in Fig7F, respectively.

## DISCUSSION

A lack of CART cell persistence limits their ability to function as an effective curative therapy [37]. It is also well documented that driving ex vivo CART expansion in the presence of abundant nutrients reduces their functional ability upon in vivo transfer [38]. While numerous studies have demonstrated that limiting nutrients to enforce starvation pathways creates metabolically optimal CARTs [4, 15, 17], multiple pathways have been highlighted, making it difficult to define the most effective intervention. Further, engineering nutrient deficient media can be costly, limiting clinical translation. Meanwhile, metabolic rewiring via promotion of mitochondrial biogenesis and FAO, while reducing ROS production, are promising interventions to improve CART function, adding yet another layer of complexity to CART manufacturing [13, 14, 19, 39]. Ultimately, the ability to generate multiple metabolic changes with one treatment could create both a greater in vivo advantage and simultaneously reduce disruptions to current CART protocols. Our data suggest that reinforcing AMPK signaling, via treatment with the direct agonist Compound 991, can achieve these goals.

Compound 991 is a commercially available agonist which binds directly to the AMPK heterotrimer [23]. We have optimized a treatment protocol which allows for maximal metabolic benefit without restricting T cell expansion, a notable limitation of other methods. T cells expanded in 991 gain mitochondrial capacity, a trait which continues upon re-stimulation and without additional agonist treatment. This gain in mitochondrial capacity correlates with increased mitochondrial density, likely occurring through mechanisms downstream of the transcription factor PGC1α [40, 41]. Indeed, overexpression of PGC1α alone has been demonstrated to improve CART therapy via its impact on mitochondrial biogenesis [12]. Impressively, despite increased mitochondrial activity in 991-treated T cells, ROS abundance was regularly reduced following restimulation, suggesting these cells also have heightened redox capacity. Indeed, AMPK is known to augment activity of the transcription factor NRF2, which drives transcriptional programs to increase redox proteins [42, 43]. Further, increased NRF2 activity alone can improve T cell metabolic function [16]. Importantly, and consistent with the greater number of 991-treated T cells recovered upon restimulation, AMPK-mediated redox buffering has been identified as a positive factor facilitating increased cellular proliferation [44]. Together, these metabolic advantages led us to hypothesize that 991 exposure during T cell expansion could improve subsequent in vivo CART cell function.

Interestingly, despite the dramatic functional improvement, we found no consistent changes in the differentiation status of 991-treated T cells, including just prior to injection. This was surprising, in part because a different model of AMPK activation promoted formation of T cell central memory populations [21]. These current studies, however, did uncover upregulation of both metabolic and cell cycle gene sets through GSEA. Coupled with the in vivo data that 991-pretreatment improves CART persistence, particularly of CD4+ cells, our results together suggest that the underlying metabolic phenotype and capacity, rather than T cell differentiation status, is the major determinant of long-term survival in our model. And while much debate has surrounded the role of memory T cells to prevent metabolic and functional exhaustion, our data support a model where metabolic and functional optimization occurs independently from changes in memory phenotype.

Even given the variability of using T cells from random human donors, the metabolic changes occurring following 991 treatment were dramatic. AMPK normally signals in the setting of nutrient starvation [20] and 991 treatment activated a metabolic program consistent with nutrient starvation despite adequate levels of nutrients in the media. This surprisingly led to reduced glucose uptake but intracellular lactate retention, with cells relying on intracellular sources of sugar carbons for the glycolysis they were pursuing. In line with intracellular nutrient utilization, agonist treatment encouraged autophagy while reducing energy expended through protein translation, adaptations implicated as beneficial for long-term cellular fitness [32]. 991-treated cells also upregulated FAO, consistent with the binding pocket for Compound 991 on AMPK being the docking site for LC-FAs to increase AMPK-driven activity [25]. Importantly, alongside increases in FAO, mitochondrial biogenesis, and autophagy, agonist treatment also simultaneously generated metabolites important for mitochondrial health and redox buffering. Indeed, glutamate, aspartate, and threonine all contribute to the generation of proline and glycine [45], the latter being products which have been highlighted in multiple models as supporting mitochondrial function and longevity [26–30]. Many of these metabolites are also critical for other metabolic pathways, including the tricarboxylic acid cycle and nucleotide synthesis [46, 47]. In fact, redirecting energy expenditure towards nucleotide synthesis during amino acid restriction has been implicated in fueling the subsequent increase in proliferative burst observed following restoration of nutrient levels [16]. Future studies to delineate AMPK’s role in directing the production of different metabolites will be important to further our specific mechanistic understanding of how the observed metabolic optimization is occurring.

With an optimized metabolic profile, we hypothesized that 991-treated CARTs would demonstrate both increased proliferative capacity and enhanced in vivo persistence. We attempted to quantify proliferation at two early times post-injection, but identified no proliferative advantage to 991-treated CARTs in either the BM or spleen. There was a trend towards increased 991-treated CART cell numbers in the bone marrow on Day 3, but this difference did not reach statistical significance. Meanwhile, we found a surprising but transient increase in DMSO-treated CART proliferation 72 hours post-injection, which was gone by one week. It is possible that leukemia clearance was slower in mice receiving DMSO-treated CART cells, allowing us to capture the final proliferative expansion of a small population of short-lived effectors cells in the spleens of these animals. In support of this interpretation, the increased proliferation at 72 hours did not translate to numerically more DMSO-treated CART cells at one week, suggesting that any actively proliferating cells at 72 hours were short-lived. The far more interesting finding became evident between days 5 and 7, where significantly more 991-treated CD4+ CART cells were recovered from both the bone marrow and spleen. These increased numbers, in the absence of proliferative advantages, strongly suggest that 991 in vitro treatment increases the resiliency of T cells at early times post-transfer, allowing them to persist longer in vivo. Further, the widening CD4/CD8 ratio, which became more dramatic over time, further implies that human CD4 T cells were the subtype most positively impacted by agonist pretreatment. Recent clinical data from long-term survivors, highlighting persistence of the CD4+ compartment as a critical factor for effective cure [48], underscores the importance of these exciting results.

Altogether, we demonstrate that expanding CARTs in the presence of the direct AMPK agonist Compound 991 metabolically reprograms them, by encouraging cellular starvation pathways without actually starving the cells and promoting FAO alongside augmentations in mitochondrial health and capacity. CARTs resulting from this process demonstrate an impressive increase in their metabolic capabilities, translating to improved in vivo persistence, particularly of donor CD4+ cells. Thus, we conclude that addition of Compound 991 to currently utilized culture methods represents an easily translatable intervention to metabolically optimize human T cells, creating products with the improved capacity to serve as an effective curative therapy.

## METHODS

### Virus Production

A CD19-targeting CAR, based on the YESCARTA protein sequence, was cloned into a pHR backbone (similar to Addgene #14858, kind gift from Jason Lohmueller, UPMC Hillman Cancer Center), followed by addition of a T2A linker and a truncated EGFR tag. Transformed bacterial cultures were grown overnight in Terrific Broth (Sigma Aldrich) and plasmids isolated using QIAGEN QIAmp Miniprep Plasmid Isolation Kit 250. HEK293Ts (ATCC) were cultured in DMEM media (Gibco #11966-025) containing 10% fetal bovine serum (FBS), Pen Strep, 2mM L-Glutamine, and MEM Non-Essential Amino Acids. Early passage cells were transfected using the Lipofectamine 3000 Transfection kit (Invitrogen) with 2500ng of RSV-REV, PMD-2G, and PRRE and 10,000ng of CAR-tEGFR plasmids. After 24 hours, supernatant was replaced with IMDM media (Gibco #12440-053) containing 10% FBS. Supernatant containing viral particles was harvested at 48 and 72 hours, combined with Lenti-Pac (GeneCopoeia), and incubated at 4 degrees C overnight. Viral supernatants were then centrifuged at 3500x (g) for 25 minutes at 4 degrees, resuspended in DMEM, and either frozen at –80C or used immediately.

### T cell isolation, transduction, and culture

De-identified buffy coats were obtained from healthy human donors (Vitalant), diluted with PBS, layered over lymphocyte separation medium (MPbio), and centrifuged at 400 xg and 25 degrees for 20 minutes with no brake. The PBMC layer was removed and T cells isolated using the Miltenyi Biotec Human Pan T cell isolation kit. Purified T cells were resuspended in AIM-V +5% SR (Gibco #A25961-01) and plated with Human T-Activator CD3/CD28 Dynabeads™ (Fisher Scientific (Thermo) 11132D) at a 2:1 ratio. For standard human T cells, cells were split on Day 3 with fresh media. For CART cell production, transduction was performed per manufacturer’s instructions utilizing retronectin coated plates (Takara) on Days 2 and 3. In both cases, cells were removed from Dynabeads by magnetic separation on Day 5 post-stimulation and expanded in AIM-V media containing 5% SR and IL-2 at 100IU/ml. Cells were re-plated with fresh media every 48 hours thereafter. Compound 991 (SelleckChem S8654, Table S1) was reconstituted in DMSO and added to cultures at a final concentration of either 10 or 25µM on Days 5 and 7. Control cultures received an equal volume of DMSO. For the survival curve and radiance analysis in Figure 3, 10uM and 25uM-treated samples were combined into one 991-treated supergroup to improve statistical power. For re-stimulation experiments, primary human T cells were re-plated with Dynabeads at a 1:1 ratio for up to 72 hours; CART cells were re-stimulated with NALM6 targets at a 3:1 ratio.

### Mice, cell lines, and xenograft leukemia model

NSG (NOD.*Cg-Prkdc^scid^Il2rgt^m1Wjl^*/SzJ) mice were purchased from Jackson Laboratories. Male and female mice were used interchangeably and housed in a specific pathogen-free facility. Recipient animals were 8-12 weeks old at the time of injection. The human NALM6 B-cell leukemia cell line was purchased from ATCC and transduced with a retroviral vector expressing Zs-Green and Luciferase, the latter a kind gift from Jason Lohmueller. NSG mice were injected with 1e6 NALM6 cells and seven days later received 3e6 total CARTs, either pre-treated with DMSO or 991. The ‘leukemia only’ control group received no CARTs. The experimental unit was a single animal. Leukemia burden was followed weekly by IVIS imaging, following intraperitoneal injection of 3 mg luciferin and imaging after 10 minutes. Any animals with a baseline radiance below 1e8 were considered leukemia-free, a metric assigned prior to the start of the experiment. One animal from the 991-treated group in Figure 3 died on day 63 while still being leukemia-free by IVIS imaging. Two bone marrow samples were lost to processing from the day 3 samples in Fig7B-D. Twenty-five NSG mice were utilized for the survival curve and radiance data presented in Fig3F-G and 43 NSG mice were used for the in vivo experiments in Figure 7. Sample size was determined based on our previous experience using these models and the number of CART cells available at the time of injection. Mice were randomly assigned to treatment groups based on the order in which they were ear-punched (also randomly assigned) and wherever possible, recipients of all three treatment groups (leukemia-only, DMSO, and 991) were co-housed prior to and following Nalm6 and CART cell injection. Cells were administered in numeric order within the cage, assuring equivalent timing between all dosing groups. Technicians performing the IVIS imaging were not aware of the treatment allocation. Survival, as the primary outcome measure, was assessed out to day 70, as determined prior to the start of the experiment.

### Protein Isolation and Immunoblot

T cells were counted, washed with PBS, and lysed with 10% trichloroacetic acid. Lysates were centrifuged at 16,000x(g) at 4*C for ten minutes, washed twice in ice cold acetone, resuspended in solubilization buffer (9M Urea containing 1% DTT and 2% Triton X and NuPAGE lithium dodecyl sulfate sample buffer 4X (Invitrogen) at a 3:1 ratio), and heated at 70*C for 10 minutes. Protein gel electrophoresis was performed on ice using NuPAGE 4-12% Bis-Tris Protein Gels (Invitrogen) at 135V. In some cases, protein samples were heated to 95C for 5 minutes prior to gel loading. Protein was transferred to Invitrolon^TM^ 0.45µm PVDF membranes (Invitrogen) at 30V on ice for one hour. Membranes were blocked in Tris Buffered Saline-Triton containing 5% nonfat milk and immunoblotting performed according to the Cell Signaling Technologies Western Blot Protocol. Blots were stripped for 10 minutes (Restore PLUS Western Blot Stripping Buffer, Thermo) prior to re-probing. Antibodies used for immunoblotting are listed in Table S2. Blots were developed with Super Signal West Femto chemiluminescence reagents (Thermo, 34096), detected by CL-X Posure Film (Thermo), and scanned in grayscale with an Epson V600 scanner. Images were cropped using ImageJ Software (version 1.47T), inverted, and densitometry quantitated in an area encompassing the largest band, followed by quantitation of subsequent bands using the same 2-dimensional area.

### Flow Cytometry

Cells were washed with PBS + 2% FBS before staining with antibodies at 1:100 dilution for 30 minutes. For intracellular stains, cells were fixed per manufacturer’s instructions using Fix/Perm kit (Invitrogen, Cat #:88-8824-00) and then stained with antibodies at 1:100 dilution. Antibodies and other flow cytometry reagents are listed in Table S3/S4. MitoTracker Green (Invitrogen) staining was performed at 50 nM in room temperature PBS for 15 minutes. CellROX (Invitrogen) staining (500 nM) was performed in culture medium for 30 minutes at 37 degrees. FAO blue (DiagnoCine Precision) staining was performed in serum-free AIMV at 15µM at 37deg for 2 hours. Nile red (Thermo) staining was performed in serum free AIMV at 0.5µg/ml at 37deg for 15 minutes. Cyto-ID (Enzo) staining was performed per manufacturer’s instructions (37 C x 30 minutes), with 500nM rapamycin added to control cultures at the time of staining. Puromycin (MedChemExpress) uptake was performed in AIMV +5%SR at 10µg/ml at 37deg for 30 minutes. In some cases, cells were pre-treated with 8µM etomoxir (Cayman Chemical Company) in AIMV + 5%SR for 15 minutes at 37deg before puromycin addition. BrdU analysis was performed utilizing the Phase-Flow kit per manufacturer’s instructions (BioLegend). *In vitro* cells were cultured in BrdU at 0.5µl per ml of cell solution per manufacturer’s instructions for 2 hours at 37deg prior to staining. Flow data was captured on a BD Fortessa analyzer (BD Biosciences) and evaluated using FlowJo software (version 10.1, Tree Star). Cells were gated by forward and side scatter to identify the lymphocyte population followed by downstream analysis.

### Seahorse Mito Stress Assay

The Seahorse XF Cell Mito Stress Test (Agilent, Santa Clara, CA; Catalog #103015-100) was run on a Seahorse XFe96 Bioanalyzer (Agilent) to determine basal and maximal oxygen consumption (OCR), spare respiratory capacity (SRC), and extracellular acidification rates (ECAR). T cells were plated in assay media (XF Base media (Agilent) with glucose (25mM), sodium pyruvate (2 mM) and L-glutamine (4 mM) (Gibco), pH 7.4 at 37 °C) on a Seahorse cell culture plate coated with Cell-Tak (Corning) at 1e5 (restim) or 1.5e5 (resting) cells/well. After adherence and equilibration, basal ECAR and OCR readings were taken for 30 min. Cells were then stimulated with oligomycin (2 µM), carbonyl cyanide 4-(trifluoromethoxy) phenylhydrazone (FCCP, 1 µM), and rotenone/antimycin A (0.5 µM) to obtain maximal respiratory and control values. Assay parameters were: 3 min mix, no wait, 3Lmin measurement, performed at baseline and repeated after each injection (3 cycles total). SRC was calculated as the difference between basal and the maximal OCR value obtained after FCCP uncoupling. The XF Mito Stress Test report generator and Agilent Seahorse analytics were used to calculate parameters using Wave software (Agilent, Version 2.6.1.53).

### RNA sequencing

Total RNA, isolated using the RNeasy Plus Mini Kit (Qiagen) in technical triplicates, was used to generate libraries using Illumina Stranded Total RNA Prep and sequenced on an Illumina Nextseq2000 at the Health Sciences Sequencing Core at the UPMC Children’s Hospital of Pittsburgh. Differentially expressed genes were generated using DEseq2, identifying genes >2-fold change in expression level and p value of 0.05 as determined by two-way ANOVA. Enrichment analysis was accomplished using GSEA software, a joint project of UC San Diego and Broad Institute [49], followed by comparison to datasets from publicly available databases.

### Metabolomics

For metabolite analysis, cells were washed and flash frozen in liquid nitrogen in technical replicates of five. Through collaboration with the University of Pittsburgh Health Sciences Mass Spectrometry Core, cells underwent metabolite extraction via resuspension in ice-cold 80% methanol, followed by addition of standards and subsequent liquid chromatography-high resolution mass spectrometry analysis. Following untargeted metabolomic analysis, putative metabolite identifications with a p value <0.05 and fold-change >2, were validated with commercial standards based on retention time, accurate mass, and MS2 fragmentation. Pathway analysis was performed on the untargeted dataset using Metaboanalyst: https://www.metaboanalyst.ca/, with comparison to the Biocyc database (Biocyc.org).

### Statistics

Graphing and statistical analysis was performed using GraphPad Prism for Windows (version 9.3.0, San Diego, CA; www.graphpad.com). Unpaired two-tailed Student t test and two-way ANOVA analysis were used to determine statistical significance. Log-rank (Mantel-Cox) analysis defined survival curve differences. Unless noted otherwise, data are displayed as mean ± standard deviation.

## Declarations

### Ethics approval and consent to participate

All animal studies were approved and carried out according to Institutional Animal Care and Use Committee guidelines from the University of Pittsburgh. All studies on human cells were given an exempt status by the University of Pittsburgh Institutional Review Board.

### Consent for publication

All authors were given a chance to review the contents of this article and have consented to its publication.

### Availability of data and material

RNA Sequencing data will be made available through a publicly accessible, online database upon publication of the manuscript. All remaining data are contained within the submitted documents. Methods and materials will be shared in accordance with standard National Institutes of Health policy by contacting the corresponding author.

### Authors’ contributions

ELB designed and performed experiments, analyzed data, generated figures, and drafted and reviewed the manuscript. MQ performed experiments and reviewed the manuscript. HS, HB, CW, and FK performed experiments. AR performed experiments and reviewed the manuscript. SM designed and performed experiments and analyzed data. AY and AP analyzed data. SG designed experiments, analyzed data, and reviewed the manuscript. CAB designed and performed experiments, analyzed data, and reviewed and edited the manuscript. Authorship order was assigned based on percent contribution to the final manuscript including overall intellectual involvement.

## Acknowledgements

Schematics created using BioRender.com

## Abbreviations

AA: Amino acid
ALL: Acute lymphoblastic leukemia
AMPK: AMP-activated protein kinase
ANOVA: Analysis of variance
BrdU: Bromodeoxyuridine
CART: Chimeric antigen receptor T cell
CPT1A: Carnitine palmitoyltransferase 1A
DMEM: Dulbecco’s modified Eagle’s media
DMSO: Dimethylsulfoxide
DTT: Dithiothreitol
ECAR: Extracellular acidification rate
EGFR: Epidermal growth factor receptor
FAO: Fatty acid oxidation
FBS: Fetal bovine serum
FCCP: Carbonyl cyanide 4-(trifluoromethoxy) phenylhydrazone
GSEA: Gene set enrichment analysis
IVIS: In vivo imaging system
LC-FA: Long-chain fatty acid
LC-HRMS: Liquid chromatography, high resolution mass spectrometry
MFI: Median fluorescence intensity
mTOR: Mammalian target of rapamycin
NSG: NOD-scid IL2Rgammanull
OCR: Oxygen consumption rate
PBMC: Peripheral blood mononuclear cell
PBS: Phosphate buffered saline
PGC1α: Peroxisome proliferator–activated receptor gamma coactivator-1 alpha
PVDF: Polyvinylidene difluoride
ROS: Reactive oxygen species
SRC: Spare respiratory capacity
ULK11: Unc-51 like kinase 1

