## Supplemental Figures for "AMPK agonism optimizes the *in vivo* persistence and anti-leukemia efficacy of chimeric antigen receptor T cells"

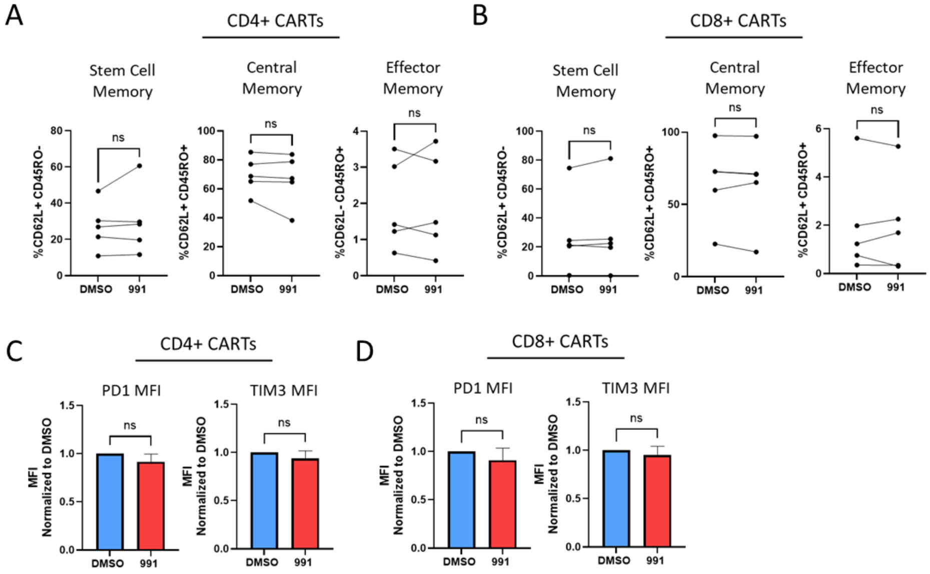


**Supplemental Figure 1. 991 exposure does not change the memory or exhaustion phenotype of CART cells.** *A-B*, Human CARTs were taken on D11 of culture and stained for memory markers CD62L and CD45RO. Data analysis was divided into CD4 (**A**) and CD8 (**B**) T cell subsets. C-D Human CARTs were recovered from culture on D11 and stained for exhaustion markers PD1 and Tim3. Data are again divided into CD4 (**C**) and CD8 (**D**) T cells.

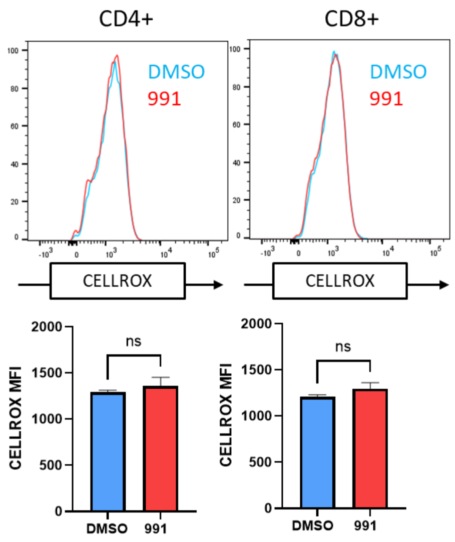


**Supplemental Figure 2. 991 treatment does not increase ROS burden during treatment.** T cells were assessed on D7 of culture for ROS burden using the ROS-reactive dye, CellROX (A). Bar graphs represent data from 2 human donors.

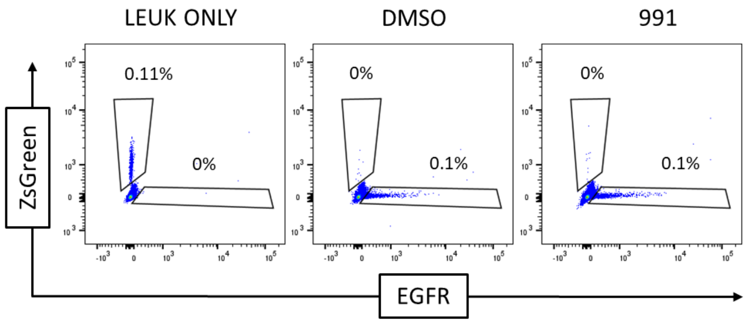


**Supplemental Figure 3. Leukemia is absent in both DMSO- and 991-treated CART recipients by Day 3 post-CART injection.** NSG mice were sacrificed 3 days after CART injection and cells of the bone marrow analyzed by flow for ongoing leukemia (using Zs-Green) and the presence of CART cells (using the CAR EGFR tag). Representative flow plots are shown for the leukemia only control, as well as the DMSO- and 991-treated CART groups.

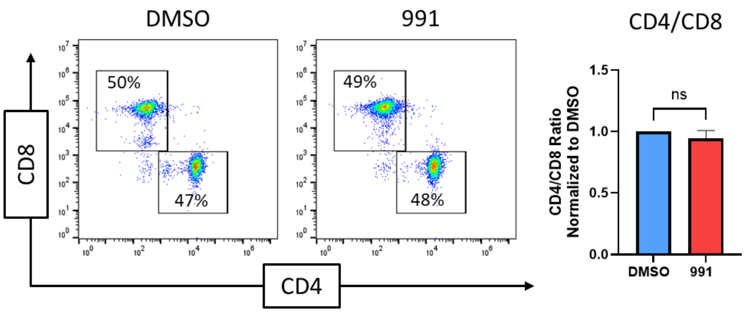


**Supplemental Figure 4. 991 exposure does not change the CD4/CD8 ratio of CART cells prior to injection.** Day 11 CART cells were stained for CD4 and CD8 and analyzed by flow. Bar graphs on the right represent composite data from 6 human donors.

**Supplemental Tables**

**Supplemental Table 1 – Reagents**

| Name | Company | Catalog # |
|---|---|---|
| **Bromodeoxyuridine (BrdU)** | *BioLegend* | 423401 |
| **BrdU kit** | *BioLegend* | 370706 |
| **D-luciferin, potassium salt** | *GOLDBIO* | LUCK-1G |
| **Etomoxir (sodium salt)** | *Cayman Chemical Company* | 11969 |
| **ex229 (compound 991)** | *Selleckchem* | S8654 |
| **Glucose Assay Kit** | *Abcam* | ab65333 |
| **L-Lactate Assay Kit** | *Sigma-Aldrich* | MAK329-1KT |
| **Puromycin dihydrochloride** | *MedChemExpress* | HY-B1743A |

**Supplemental Table 2 – Antibodies for Immunoblot analysis**

| Antigen | Company | Clone | Catalog # |
|---|---|---|---|
| **Phospho-ULK-1** | *Cell Signaling* | D1H4 | 5869S |
| **ULK-1** | *Cell Signaling* | D8H5 | 8054S |
| **Phospho-AMPK** | *Cell Signaling* | T172 | 2535S |
| **AMPK** | *Cell Signaling* | F6 | 2793S |
| **Beta Actin** | *Cell Signaling* | 13E5 | 4970S |
| **PGC1α** | *NOVUS* | (polyclonal) | NBP1-04676 |
| **CPT1A** | *Cell Signaling* | D3B3 | 12252S |
| **Phospho-Raptor** | *Cell Signaling* | E4V6C | 89146S |
| **Raptor** | *Cell Signaling* | 24C12 | 2280S |

Abbreviations: ACC Acetyl CoA Carboxylase, AMPK AMP-activated protein kinase, ULK1 Unc51-like kinase 1, PGC1α PPARG coactivator 1 alpha, CPT1A carnitine palmitoyltransferase 1A, Raptor regulatory associated protein of mTOR

**Supplemental Table 3 – Antibodies for flow cytometry**

| Antigen | Company | Clone | Conjugate | Catalog # |
|---|---|---|---|---|
| **BrdU** | *BioLegend* | 3D4 | AF647 | 364108 |
| **CD4** | *BioLegend* | SK3 | Pac Blue | 344620 |
| **CD4** | *Invitrogen* | RPA-T4 | PeCY7, APC, PE | 17-0049-42 |
| **CD8** | *Invitrogen* | SK1 | APCeF780, PerCPAeF710 | 47-0087-42 |
| **CD8** | *BioLegend* | SK1 | BV711 | 344734 |
| **CD45RO** | *BD Biosciences* | UCHL | BV711 | 563722 |
| **CD62L** | *Invitrogen* | DREG-56 | PE | 12-0629-41 |
| **EGFR** | *BioLegend* | Ay12 | PE, PECy7 | 352904 |
| **PD-1 (CD279)** | *BioLegend* | NAT105 | AF488 | 367408 |
| **Puromycin** | *BioLegend* | 2A4 | AF647, PE | 381504 |
| **TIM3** | *BioLegend* | F38-2E2 | APC | 345011 |

**Supplemental Table 4 – Reagents for flow cytometry**

| Name | Company | Catalog # |
|---|---|---|
| **LIVE/DEAD™ Aqua** | *Invitrogen* | L34957 |
| **MitoTracker Green FM** | *Invitrogen* | M7514 |
| **CellROX Deep Red** | *Invitrogen* | C10491 |
| **Nile Red** | *Invitrogen* | N1142 |
| **FAO Blue** | *Diagnocine Precision* | FNK-FDV-0033 |
| **CYTO-ID** | *Enzo* | ENZ-51031-0050 |
